# Umbilical cord blood derived microglia-like cells to model COVID-19 exposure

**DOI:** 10.1101/2020.10.07.329748

**Authors:** Steven D. Sheridan, Jessica M. Thanos, Rose M. De Guzman, Liam T. McCrea, Joy Horng, Ting Fu, Carl M. Sellgren, Roy H. Perlis, Andrea G. Edlow

## Abstract

Microglia, the resident brain immune cells, play a critical role in normal brain development, and are impacted by the intrauterine environment, including maternal immune activation and inflammatory exposures. The COVID-19 pandemic presents a potential developmental immune challenge to the fetal brain, in the setting of maternal SARS-CoV-2 infection with its attendant potential for cytokine production and, in severe cases, cytokine storming. There is currently no biomarker or model for *in utero* microglial priming and function that might aid in identifying the neonates and children most vulnerable to neurodevelopmental morbidity, as microglia remain inaccessible in fetal life and after birth. This study aimed to generate patient-derived microglial-like cell models unique to each neonate from reprogrammed umbilical cord blood mononuclear cells, adapting and extending a novel methodology previously validated for adult peripheral blood mononuclear cells. We demonstrate that umbilical cord blood mononuclear cells can be used to create microglial-like cell models morphologically and functionally similar to microglia observed *in vivo*. We illustrate the application of this approach by generating microglia from cells exposed and unexposed to maternal SARS-CoV-2 infection. Our ability to create personalized neonatal models of fetal brain immune programming enables non-invasive insights into fetal brain development and potential childhood neurodevelopmental vulnerabilities for a range of maternal exposures, including COVID-19.

## Introduction

Maternal immune activation can result from exposures ranging from metabolic conditions, to stress and infection, with potential *in utero* consequences to the developing fetus. ^1-6^ In particular, epidemiologic studies strongly suggest that maternal viral and bacterial infections in pregnancy may be associated with adverse neurodevelopmental outcomes in offspring, particularly autism spectrum disorders, schizophrenia, and cerebral palsy, but potentially including mood and anxiety disorders as well.^1-3, 7-9^ For instance, individuals who were fetuses during the 1957 influenza pandemic had a significantly increased risk for being hospitalized for schizophrenia as an adult.^10^ The magnitude of these effects varies, but their consistency is difficult to ignore. Although mechanisms underlying neurodevelopmental morbidity in offspring remain unclear, microglial priming toward a pro-inflammatory phenotype with consequent altered synaptic pruning has been suggested as a candidate mechanism.^11-17^

Microglia, brain-resident tissue macrophages, play a key role in normal neurodevelopment by modulating synaptic pruning, neurogenesis, phagocytosis of apoptotic cells, and regulation of synaptic plasticity.^18-21^ Fetal yolk sac-derived macrophages are the progenitors for the permanent pool of brain microglia throughout an individual’s lifetime.^22-25^ As such, inappropriate fetal microglial priming (“trained immunity” ^26^) due to *in utero* immune activation may have lifelong neurodevelopmental consequences. The central role of mononuclear cells, including macrophages, in COVID-19 pathogenesis^27^ suggests that the potential risk to exposed fetal microglia requires investigation.

We have previously developed and validated adult patient-specific models of microglia-mediated pruning by reprogramming induced microglial cells from human peripheral blood mononuclear cells (PBMCs), and assaying them with isolated synapses (synaptosomes) derived from neural cultures differentiated from induced pluripotent stem cells (iPSCs).^28, 29^ We demonstrated robust evidence of abnormalities in microglia and synaptosomes from individuals with schizophrenia, shown to be complement-dependent through a C3 receptor neutralizing antibody and rescued in a dose-responsive fashion with a small molecule probe.^28^

In this study, we investigated whether our validated reprogramming methods for adult PBMCs could be successfully adapted and applied to umbilical cord blood-derived mononuclear cells (CB-MNCs) from both SARS-CoV-2 infected and uninfected pregnancies, to create personalized models of fetal brain microglia. Such models could have a wide range of application in investigating effects of *in utero* exposure on neurodevelopment. To illustrate this application, we demonstrate successful induction of microglia-like cells (CB-iMGs) from CB-MNCs from both infected and uninfected pregnancies.

## Materials/Subjects and Methods

### Ethical statement

This study was approved by the Partners Institutional Review Board. Informed consent was obtained from all participants.

### Isolation and preparation of mononuclear cells from umbilical cord blood (CB-MNCs)

Umbilical cord blood was collected at the time of delivery into EDTA tubes for plasma and CB-MNC isolation. After spinning at 1000*g* for 10 minutes to separate plasma, samples were processed for CB-MNC isolation using a Ficoll density gradient.^30^ Briefly, blood was transferred into a 50mL conical tube and then diluted to 1:1 ratio with Hanks’ Balanced Salt Solution without calcium or magnesium (HBSS minus). This diluted blood was then gently layered on top of Ficoll at 2:1 ratio (2 volumes of blood diluted with HBSS minus to 1 volume Ficoll). The conical tube was then centrifuged at 1000 g for 30 minutes at room temperature with brake inactivated to allow layering of cellular components. The cloudy ring below the plasma and above the Ficoll (i.e. the CB-MNC layer) was collected and placed in a new 15mL conical tube, with HBSS minus added to bring the volume to 15 mL. This tube was then centrifuged at 330 g for 10 minutes with high brake. The supernatant was removed and the CB-MNC pellet was washed with HBSS minus and resuspended in 10 mL HBSS minus for counting. Cells were counted on a hemocytometer in a 1:10 dilution of trypan blue. Cells were frozen in freezing medium consisting of RPMI 1640 Medium with 1% penicillin-streptomycin, L-glutamine, 1% sodium pyruvate, 1% non-essential amino-acids, 20% Fetal Bovine Serum (FBS) and 10% DMSO at 5-10 million cells/vial, placed in a chilled Mr. Frosty, then into −80 C. The following day, CB-MNC cryovials were transferred to liquid nitrogen for long-term storage. Isolated cryopreserved adult peripheral blood mononuclear cells were obtained from a single healthy control donor by Vitrologic (https://vitrologic.com) cat# MNC-300.

### Derivation of induced microglia-like cells from CB-MNCs by direct cytokine reprogramming

iMGs were derived using previously described methods^28, 29^ while CB-iMGs were derived from CB-MNCs with modifications as noted. Briefly, cryopreserved PBMCs or CB-MNCs were rapidly thawed at 37°C, diluted into media consisting of RPMI 1640 with 10% FBS and 1% Penicillin/Streptomycin. The cell suspension was centrifuged at 300g for 5 minutes at room temperature, with the brake off. After aspirating the supernatant, the cell pellet was resuspended in media, counted and plated on Geltrex-coated 24-well plates at 1⨯10^6^ cells per 0.5 mL per well. After cells were incubated at 37°C for 24 hours, the media was completely replaced with RPMI 1640 with GlutaMAX, 1% Penicillin-Streptomycin, 100 ng/ml of human recombinant IL-34 (Peprotech) and 10 ng/ml of GM-CSF (Peprotech). Media was replaced on day 13 after the initial cytokine reprogramming, and real-time live cell imaging or immunocytochemistry was performed on day 14.

### iPSC generation and neural differentiation for synaptosome isolation

iPSCs were reprogrammed from fibroblasts and used to derive neural progenitor cells, which were differentiated into neural cultures, as previously described.^28, 29^ In brief, adult human fibroblasts were reprogrammed to iPSCs using non-integrating synthetic RNA pluripotency factors, expanded and cryopreserved by Cellular Reprogramming, Inc. (https://www.cellular-reprogramming.com). To initiate neural progenitor induction, iPSCs were cultured feeder-free in E8 medium (Gibco) on Geltrex coated six-well plates and passaged using 50 mM EDTA and trituration with ROCK inhibitor (10 mM Thiazovivin; Stemgent). iPSCs were further purified using magnetic-activated cell sorting with Tra-1-60 microbeads (Miltenyi Biotec) on LS columns as described by vendor. Neural progenitor cells (NPCs) were derived from these iPSCs using Neurobasal Medium (Thermo Fisher Scientific) with 1X Neural Induction Supplement (Thermo Fisher Scientific), expanded using a neural expansion medium, and purified by double sorting using MACS against CD271 and CD133. NPCs were immunostained for markers, including Nestin, SOX1, SOX2, and Pax-6. Validated NPCs were seeded for neural differentiation on Geltrex-coated T1000 5-layer cell culture flasks (Millipore Sigma # PFHYS1008) and grown in neuronal differentiation medium (Neurobasal media (Gibco # 21103049) supplemented with 1X each (N2 supplement (Stemcell Technologies SCT # 7156), B27 supplement without Vitamin A (Gibco #12587010), Non-essential amino acids (NEAA Gibco # 11140050), penn/strep), 1 uM Ascorbic Acid, 10ng/mL BDNF and GDNF (Peprotech) and 1ug/ml mouse laminin (Sigma # L2020) for 8 weeks.

### Synaptosome isolation by sucrose gradient

Synaptosome isolation by sucrose gradient was adapted for iPSC-derived differentiated neural cultures from previously described protocols.^31-33^ First, media was aspirated from flasks and cells were washed or scraped with 1X gradient buffer (ice-cold 0.32M sucrose, 600mg/L Tris, 1 mM NaH_3_CO_3,_ 1mM EDTA, pH 7.4 with added HALT protease inhibitor - ThermoFisher # 78442), homogenized using a dounce homogenizer and centrifuged at 700 *g* for 10 minutes at 4°C. The pellet was resuspended in 10ml of 1X gradient buffer (transferred to a 30 mL and centrifuged at 15,000 *g* for 15 minutes at 4°C. The second pellet was resuspended in 1X gradient buffer and slowly added on top of a sucrose gradient with 1X gradient buffer containing 1.2M (bottom), 0.85M (middle) sucrose layers. The gradient and cell mixture was centrifuged at 26,500 rpm (∼80,000 *g*) for 2 hours at 4°C, with the brake set to “slow” so as not to disrupt the final bands. The mixtures were handled carefully and bands were inspected to confirm successful fractionation. The synaptosome band (in between 0.85M and 1.2M sucrose) was removed with a 5-mL syringe and 19Gx1 1/2” needle and centrifuged at 20,000 *g* for 20 minutes at 4°C. The final pellet was resuspended in an appropriate volume of 1X gradient buffer with 1mg/mL bovine serum albumin (BSA) with protease and phosphatase inhibitors, aliquoted and slowly frozen at - 80°C. Protein concentration was measured by BCA, and synaptosomes were analyzed by transmission electron microscopy, and immunoblotted for synaptic markers.

### Real-time live cell imaging of synaptosome phagocytosis by CB-iMGs

Real-time live cell imaging of CB-iMGs was performed as previously described.^28, 29^ Briefly, cells were imaged on the IncuCyte ZOOM live imaging system (Essen Biosciences) while incubated at 37°C with 5% CO_2_. Synaptosomes were sonicated and labeled with pHrodo Red SE (Thermo Fisher Scientific) and added to CB-iMGs at 15 µg total protein per well in 24-well plates. Phase-contrast and red fluorescence channel images were taken at a resolution of 0.61 µm per pixel every 45 minutes for a total of 315 minutes. Images were exported as 16-bit grayscale files and analyzed using CellProfiler^34^ to quantify cells and phagocytized particles. CellProfiler pipeline description and files are included in Supplemental Materials.

### Immunocytochemistry and confocal microscopy

iMGs were fixed with 4% paraformaldehyde in PBS for 15 minutes at 4°C, washed twice with PBS, blocked for 1 hour with 5% FBS and 0.3% Triton-X (Sigma Aldrich) in PBS, then washed 3 times with 1% FBS in PBS. Cells were incubated with primary antibodies in 5% FBS and 0.1% Triton-X overnight at 4°C (Anti-IBA1, 1:500, Abcam #ab5076; Anti-CX3CR1, 1:100, Abcam ab8021; Anti-PU.1, 1:1,1000, Abcam #ab183327, and Anti-P2RY12, 1:100, Alomone Labs). Cells were then washed twice with 1% FBS in PBS and incubated in secondary antibodies (1:500) and Hoechst 33342 (1:5,000) in 5% FBS and 0.1% Triton-X in PBS for 45 minutes at 4°C, light-protected. Cells were washed twice and imaged using the IN Cell Analyzer 6000 (Cytiva).

## Results

### Generating and characterizing human microglia-like cells from umbilical cord blood-derived mononuclear cells (CB-iMGs)

We adapted previously-reported methods^28, 29^ for generating iMGs from adult-derived PBMCs to reprogram umbilical cord-derived mononuclear cells (CB-MNCs) from neonates of SARS-CoV-2 negative (n=4) and SARS-CoV-2 positive (n=2) mothers, delivered between June 7, 2020 and July 6, 2020. After 2 weeks of cytokine exposure, analogous to PBMC-derived iMGs, CB-iMGs displayed typical ramified microglial morphology (Fig. 1a, i) and stained positive for canonical microglial markers IBA1, CX3CR1, PU.1, and P2RY12 (Figs. 1a, ii-v), demonstrating that iMGs can be generated from umbilical cord blood-derived CB-MNCs. These cells exhibit morphologic and features and markers of cell identity comparable to those we have demonstrated for adult blood-derived PBMCs (Figs. 1b, i-v).

**Figure 1.**
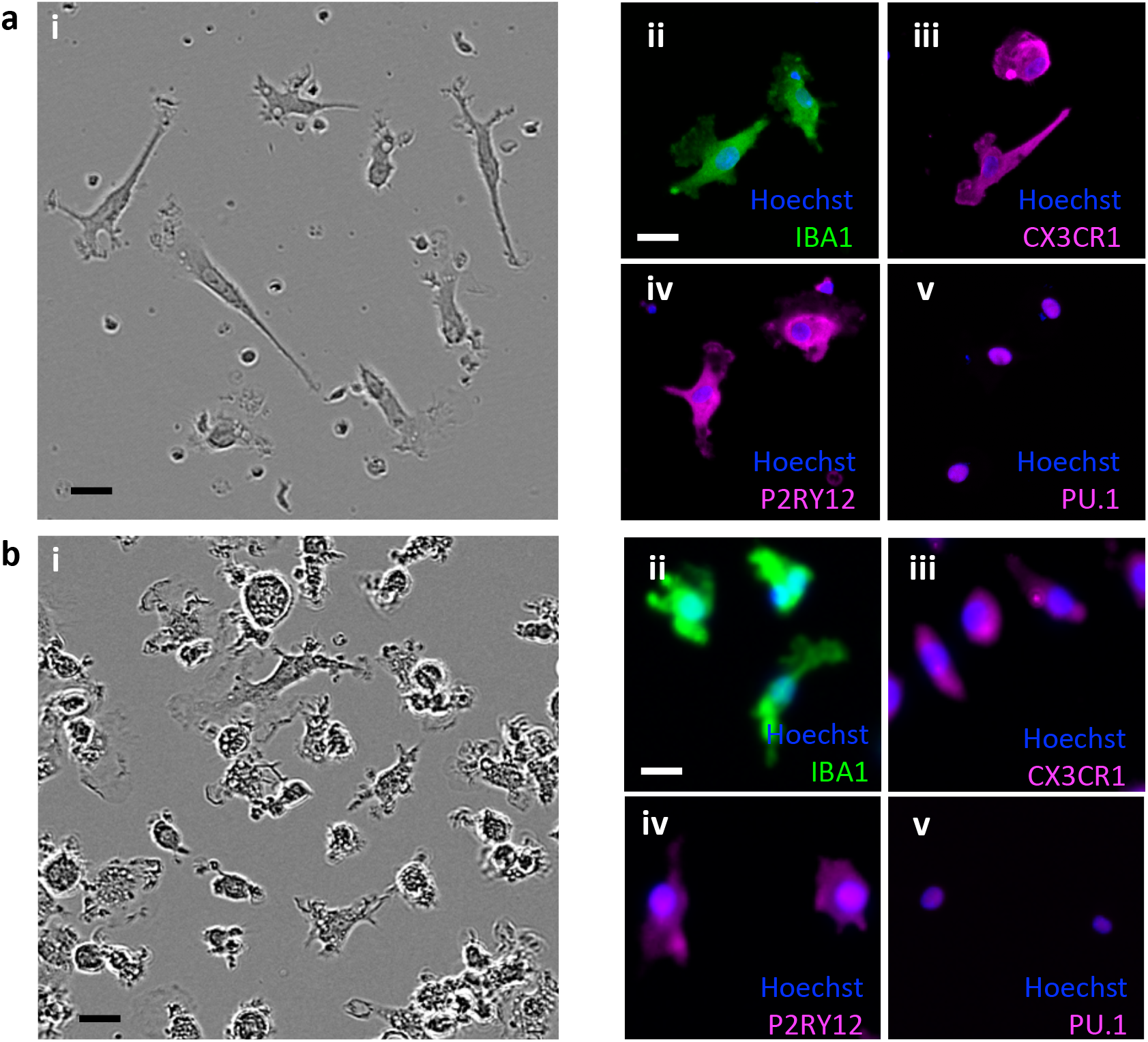
Characterization of monocyte-derived induced microglia-like cells (iMGs) by direct cytokine reprogramming. (a) Umbilical cord monocyte-derived CB-iMGs. (b) Adult PBMC-derived iMGs. (i) Morphology by phase contrast; immunostained images of iMG cells stained with nuclei (Hoechst) and indicated microglial markers (ii) IBA1, (iii) CX3CR1, (iv) P2RY12 and (v) PU.1. Scale bar: 30 μm.

### CB-iMGs demonstrate capacity to engulf isolated synaptic material in an in-vitro model of synaptic pruning

We next characterized CB-iMG function in a model of synaptic pruning using highly purified isolated nerve terminals (synaptosomes), allowing quantitation of synaptic engulfment *in vitro* with greater signal-to-noise than intact neural cultures. ^28, 29^ Figure 2a illustrates the real-time imaging-based phagocytosis assay workflow with CB-iMG cultures upon addition of iPSC-derived neural culture purified synaptosomes labelled with pHrodo Red SE, a pH-sensitive dye that fluoresces upon localization to lysosomes post-engulfment. Engulfment of synaptosomes can be robustly quantified in real-time live imaging (Figs. 2b and c) as well as by endpoint confocal microscopy of fixed immunostained cells.^28, 29^

**Figure 2.**
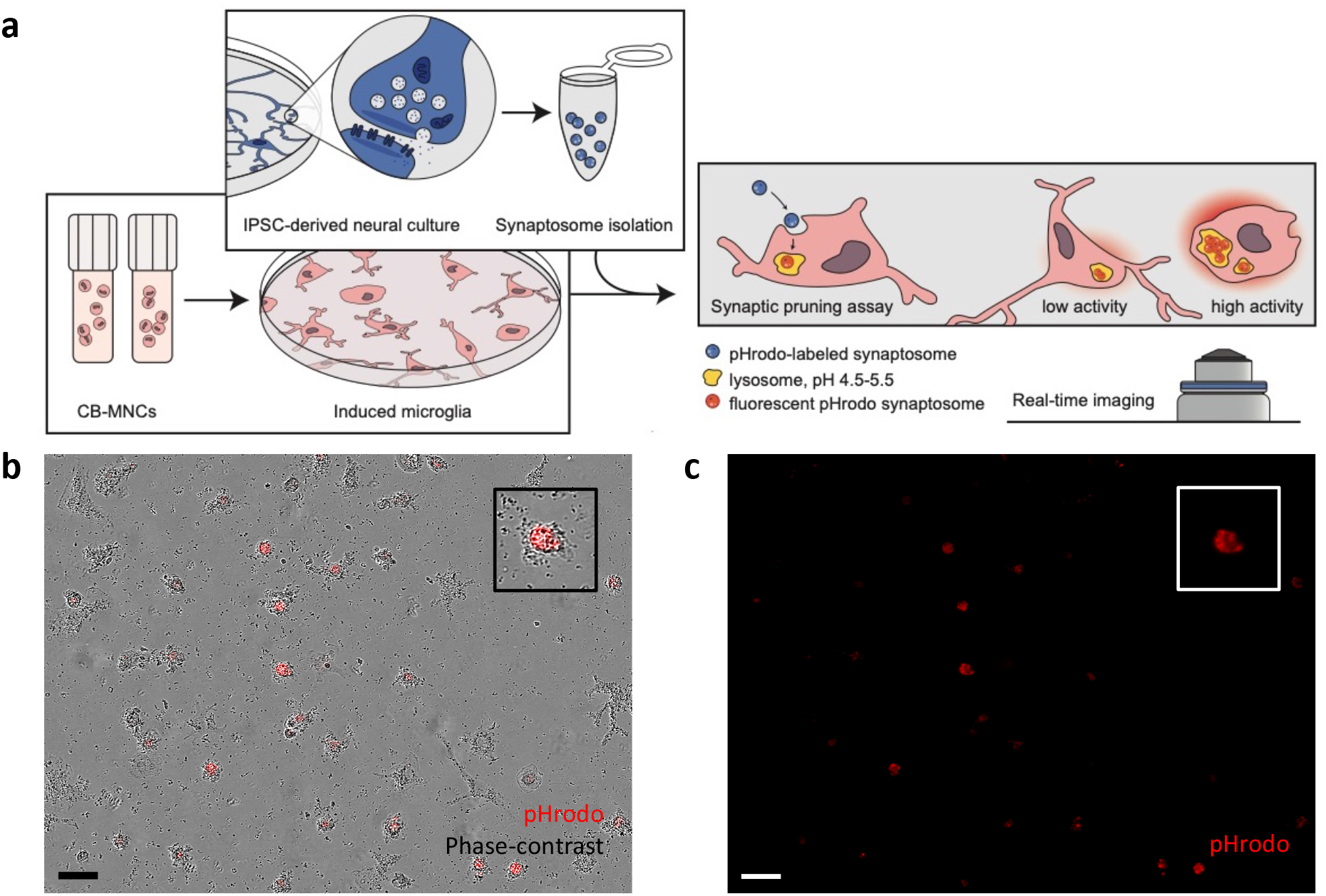
Synaptosome engulfment functional characterization of CB-iMGs in an in-vitro model of synaptic pruning. (a) Overall schematic of pHrodo-labelled quantitative synaptosome phagocytosis assay by CB-iMGs. (b) Representative live real-time images in phase-contrast/red fluorescence overlay mode showing cellular uptake and (c) red fluorescence channel alone of pHrodo (red)-labeled synaptosomes uptake after 5 h. Scale bar: 60 μm (Boxes show magnified view of engulfing CB-iMG).

To characterize kinetics of phagocytosis over time, pHrodo labeled-synaptosomes were added to the CB-iMG cultures and engulfment was visualized every 45 minutes for approximately 5 hours using real-time live fluorescence imaging (Figure 3a). Phagocytotic index was determined at each time point by measuring pHrodo area per cell (Figure 3b) using CellProfiler as described in Methods and Supplemental Information. CB-iMGs demonstrate robust phagocytosis of synaptosomes, with a time course of synaptosome engulfment rising over time qualitatively similar to that observed with adult PBMC-derived iMGs. ^28, 29^

**Figure 3.**
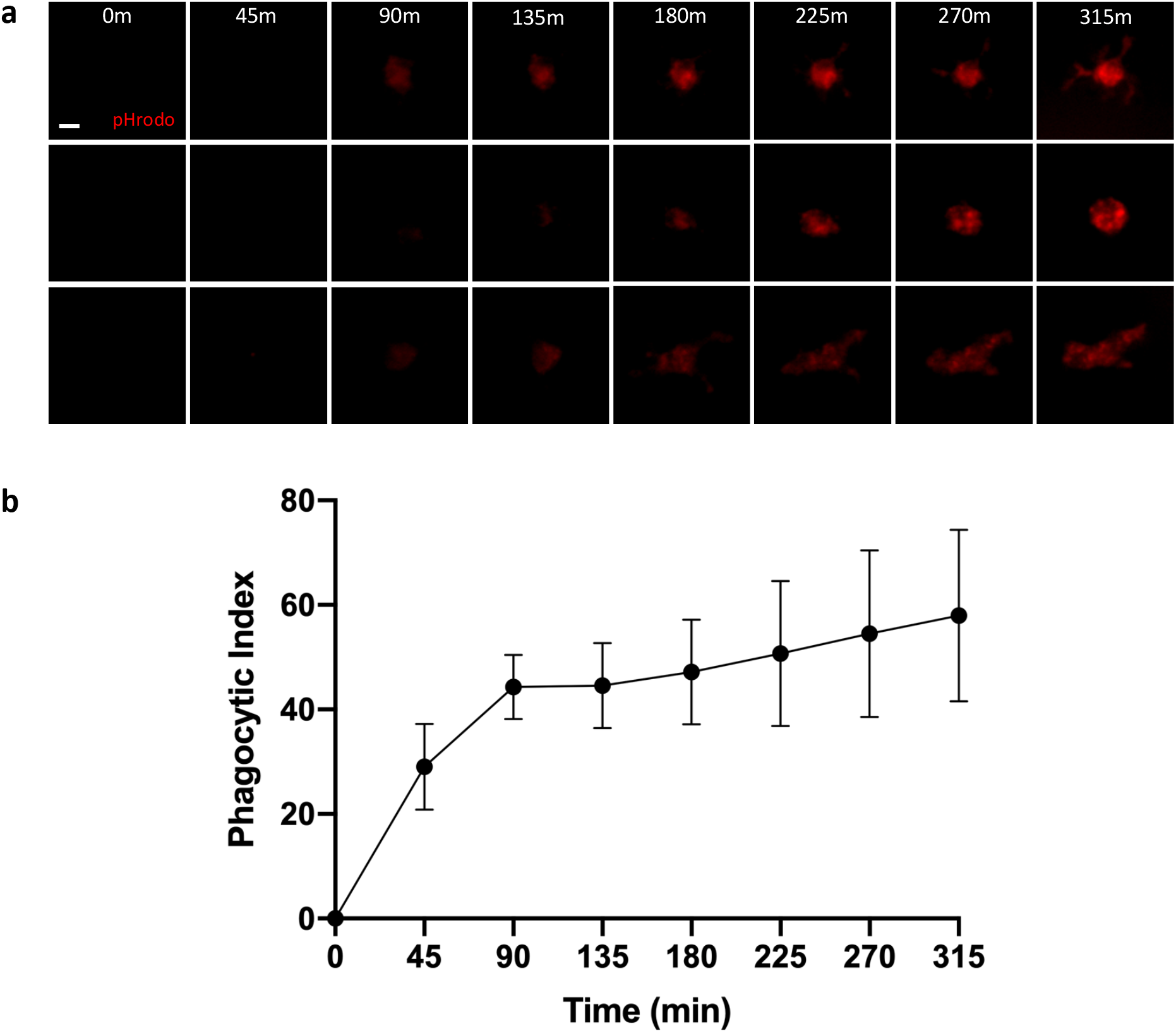
Functional Characterization of synaptosome engulfment by CB-iMGs (a) Representative pHrodo (red)-labeled synaptosome engulfment in iMG cells during live real-time imaging used for quantification Scale bar: 20 μm (b) Quantification of labeled synaptosome uptake by CB-iMGs cells during live imaging. The phagocytic index represents the mean pHrodo+ area per iMG cell over N=8 fields per well x 3 wells per line and N=4 separate healthy control CB-IMG line derivations. Error bars represent SEM.

### CB-IMGs derived from CB-MNCs exposed to maternal SARS-CoV-2 infection

To illustrate the application of these models to study maternal exposures, we generated CB-iMGs from umbilical cord blood of neonates from mothers who tested positive for SARS-CoV-2 (n=2). As observed above for CB-iMGs derived from SARS-CoV-2 unexposed pregnancies, these cells also display ramified and amoeboid morphology by phase-contrast imaging (Fig. 4a), express microglial-specific markers IBA1, CX3CR1, PU.1 and P2RY12 by immunostaining (Fig. 4b, i-iv), and engulf pHrodo-labeled synaptosomes in our synaptic pruning assay as visualized by real-time live imaging (Figs. 5a-c) and quantification (Fig. 5d). These results demonstrate the capacity to create phenotypically characteristic and functionally active CB-iMGs in a quantitative model of synaptic pruning to provide for further investigations into potential effects of maternal exposures such as SARS-CoV-2.

**Figure 4.**
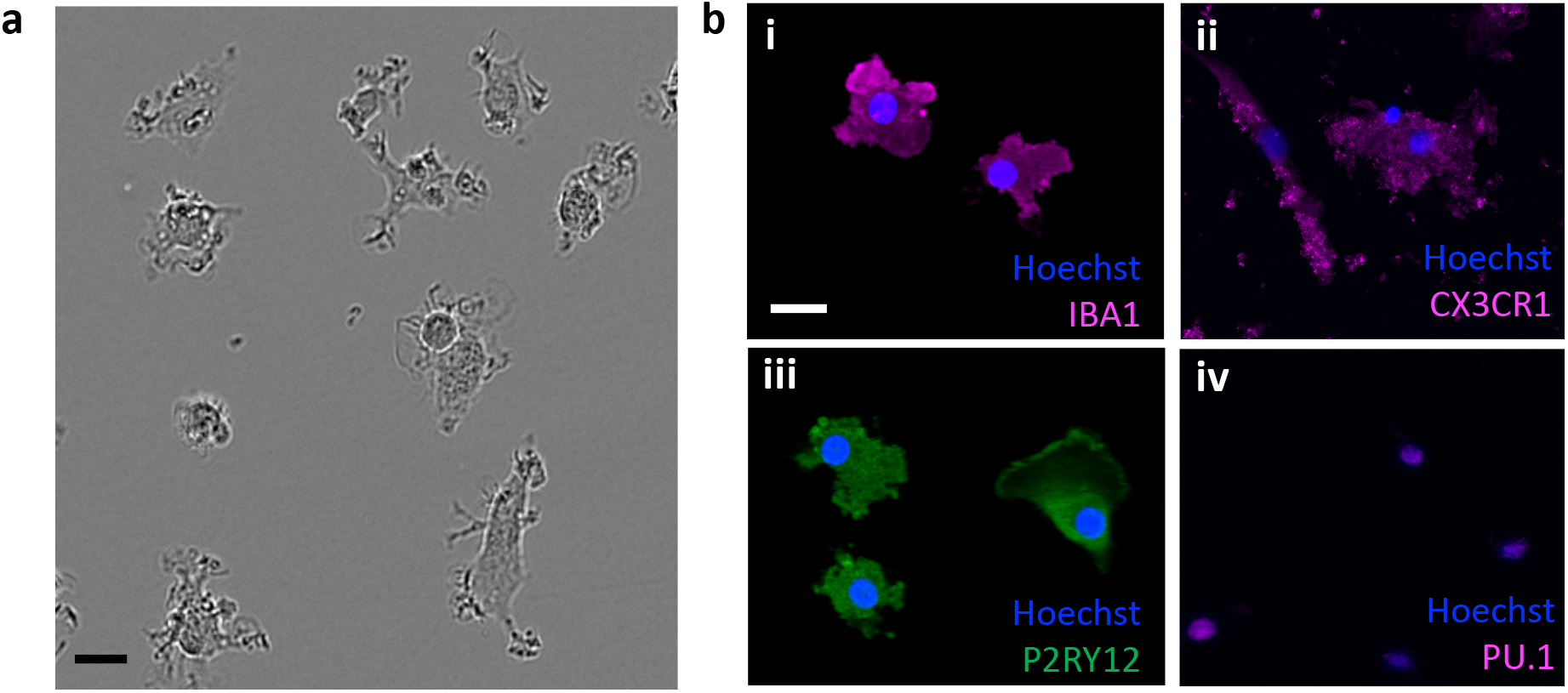
Phenotypic characterization of CB-iMGs derived from CB-MNCs exposed to maternal SARS-CoV-2 infection. (a) Phase contrast ramified morphology of patient-derived CB-iMGs, Scale bar: 30 μm. (b) immunostained images of exposed CB-iMG cells with nuclei (Hoechst) and indicated microglial markers (i) IBA1, (ii) CX3CR1, (iii) P2RY12 and (iv) PU.1. Scale bar: 25 μm.

**Figure 5.**
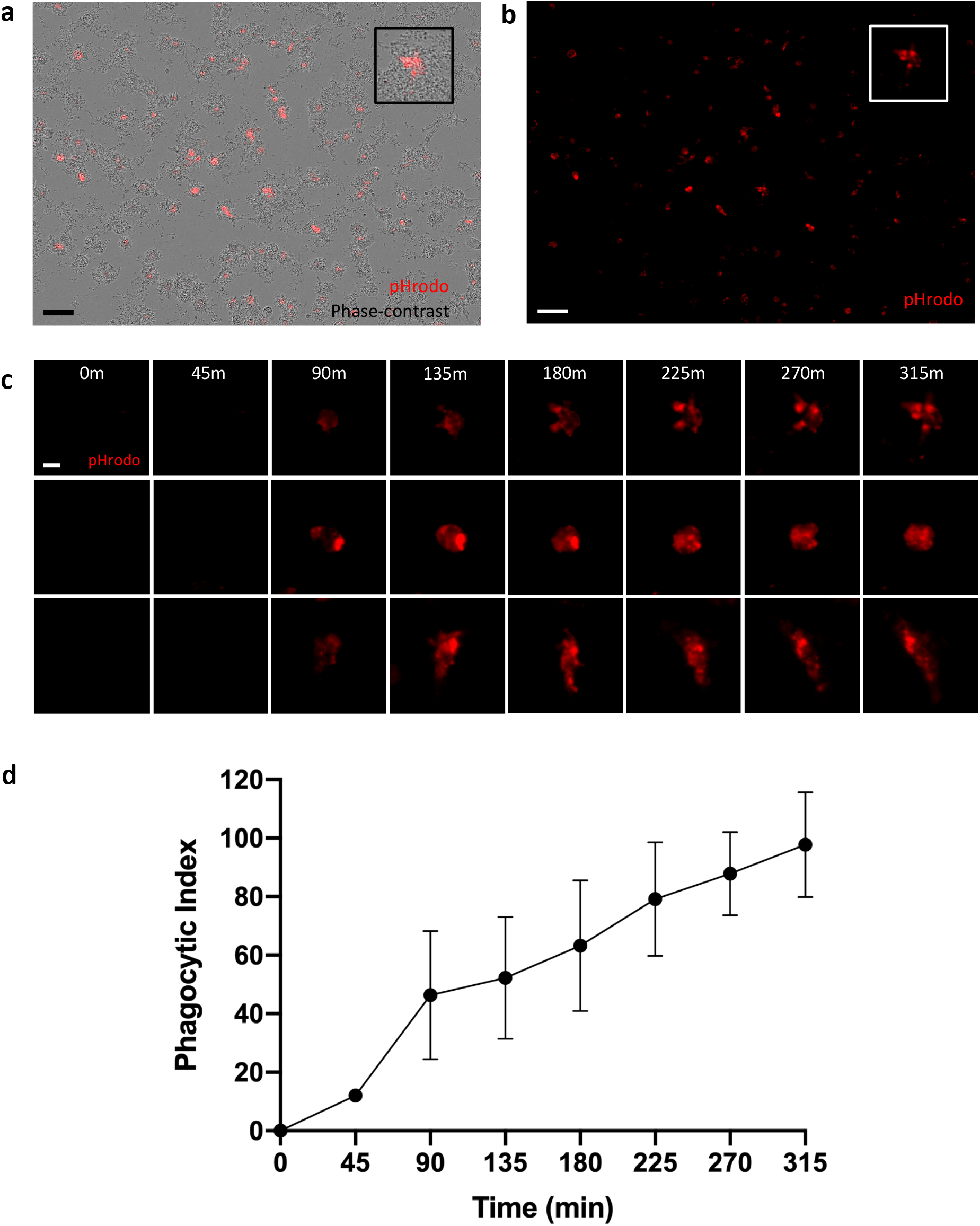
Functional characterization of synaptosome engulfment by CB-iMGs derived from CB-MNCs exposed to maternal SARS-CoV-2 infection. (a) Representative live real-time images in phase-contrast/red fluorescence overlay mode showing cellular uptake and (b) red fluorescence channel alone of pHrodo (red)-labeled synaptosomes uptake after 5 h. Scale bar: 60 μm. (Boxes show magnified view of engulfing CB-iMG) (c) Representative pHrodo (red)-labeled synaptosome engulfment in exposed CB-iMGs during live real-time imaging used for quantification. Scale bar: 20 μm. (d) Quantification of labeled synaptosome uptake by exposed CB-iMGs cells during live imaging. The phagocytic index represents the mean pHrodo+ area per iMG cell over N=8 fields per well x 3 wells per line and N=2 separate SARS-CoV-2 exposed CB-IMG line derivations. Error bars represent SEM.

## Discussion

Our results demonstrate the successful creation of neonatal patient-specific models of microglia-mediated synaptic pruning via cellular reprogramming of neonatal cord blood mononuclear cells. These models should facilitate novel insights into fetal brain development in the setting of maternal exposures, including but not limited to SARS-CoV-2 infection. Both SARS-CoV-2-exposed and unexposed umbilical cord blood derived microglia-like cells express canonical microglial markers IBA1, CX3CR1, PU.1, and P2RY12, and demonstrate a range of morphologies with varying degrees of ramification, potentially reflecting a range of activation states that can be perturbed in experimental systems. Importantly, the induced microglia phagocytose synaptosomes demonstrating that they can recapitulate this key function of microglia in the developing brain. This work suggests the potential for umbilical blood mononuclear cells to serve as a non-invasive, personalized biomarker of fetal brain microglial priming. To our knowledge, microglia have not previously been modeled from umbilical cord blood or used to predict neurodevelopmental vulnerability at a time when there is a window for intervention. These models provide the potential for quantifiable endpoints that can be used to assess microglial programming in the setting of various maternal exposures, and later can be used to test the efficacy of potential therapies to ameliorate in utero priming of microglia toward a pro-inflammatory phenotype.^35^

Previous studies have demonstrated the impact of various maternal exposures, including maternal stress, metabolic disorders, air pollution, and infections, on fetal brain development.^2-4, 7^ Such exposures may lead to offspring neurodevelopmental morbidity via maternal immune activation,^36-43^ which results in aberrant microglial programming in the developing brain.^12-15, 18^ In turn, maternal immune activation models have pointed to aberrant differentiation of fetal microglia and dysregulation of cytokine networks as key mechanisms underlying abnormal fetal brain development, with microglia primed toward a pro-inflammatory phenotype and altered synaptic pruning implicated in offspring morbidity.^11-15^ Given the extent of synapse formation and pruning that occurs in fetal and neonatal life, developmental microglial function represents a critical target for investigation.

We have previously demonstrated feasibility for the concept of using other monocyte populations to model microglial behavior using another monocyte type, fetal placental macrophages or Hofbauer cells, as a potential biologic surrogate for fetal microglial function in pre-clinical models of maternal obesity,^16^ and have shown that maternal obesity-associated inflammation primes both fetal brain microglia and resident placental macrophages toward a highly correlated pro-inflammatory phenotype.^16^ Personalized assays using more readily available cord blood mononuclear cells, as presented here, can be used to assess both baseline microglial function, and behavior in response to “second hit” inflammatory stimuli, which will be helpful in informing risk assessments for an individual fetus.

The COVID-19 pandemic, with its associated maternal immune activation and pro-inflammatory cytokine-mediated physiology,^44, 45^ may pose risk to the developing fetal brain. While current data suggest that vertical transmission of SARS-CoV-2 is relatively rare,^46-48^ the profound immune activation observed in a subset of infected individuals suggests that, even if the virus itself does not cross placenta, the developing fetal brain may be impacted by maternal inflammation and altered cytokine expression during key developmental windows.^49-51^ In this work, we suggest a model system that may be applied to investigate risk associated with the COVID-19 pandemic. Extending our work with adult peripheral blood mononuclear cells to cord blood-derived microglial models will allow for rapid, scalable models to investigate risk, yielding non-invasive, personalized assays of the impact of SARS-CoV-2 on fetal brain microglial priming and synaptic pruning function. This approach can complement more traditional approaches that will require large longitudinal cohort studies and may require years or even decades (in the case of schizophrenia, for example) to fully capture risk. The ability to detect priming of fetal brain microglia toward a proinflammatory phenotype extends beyond SARS-CoV-2, to include numerous other maternal infections in pregnancy, as well as the myriad maternal exposures that have been suggested to impact fetal microglial development.

In sum, we demonstrate the potential for umbilical cord blood mononuclear cells to serve as a non-invasive, personalized model of fetal brain microglial priming. These models provide the potential for quantifiable endpoints that can be used to assess microglial programming in the setting of various maternal exposures. We determined that CB-iMGs can recapitulate microglial characteristics and function *in vitro*, providing key insight into cells from the neonatal brain that are otherwise inaccessible at birth and throughout childhood. We illustrated their application to investigate effects of maternal infection, including SARS-CoV-2, on the developing brain.^28^ Beyond characterizing any consequent abnormalities, the scalability of this approach may enable investigation of targeted therapeutic strategies to rescue such dysfunction.

## Supporting information

Supplmental CellProfiler Analysis Pipelines

## Acknowledgements

We are grateful Dana Cvrk, C.N.M., Muriel Schwinn, N.P; Lydia Shook, M.D., Adeline Boatin, M.D., M.P.H.; Robin Azevedo, R.N.; Laurel Gardner, R.N.; Suzanne Stanton, R.N.; Natalie Croul, B.A.; Nicola Young, B.A.; Samantha Devane, B.S. for their assistance with recruitment and sample collection; to all members of the MGH Obstetric-Pediatric COVID-19 Biorepository Processing Team for their assistance with sample processing and storage, to Lael Yonker, M.D., and Alessio Fasano, M.D. for their partnership on the Obstetric-Pediatric biorepository; to Jon Li, M.D. and Xu Yu, M.D. for critical infrastructural and regulatory support; and to Anjali Kaimal, M.D.,M.A.S., and Jeffrey Ecker, M.D., for assistance with study infrastructure and departmental support. Most importantly, we thank the participants for being part of the study.

This work was supported by R01-MH120227 (Dr. Perlis) and R01HD100022 and 3R01HD100022-02S2 (Dr. Edlow).

## CellProfiler image analysis pipeline description for attached Supplemental pipeline files

CellProfiler (version 3.1.9)1 was used to measure cell counts and synaptosome area using phase-contrast and red fluorescence live-cell images, respectively. Baseline phase-contrast images were processed using the EnhanceOrSuppressFeatures module. RobustBackground method thresholding was applied to processed images and used to identify cells based on size using the IdentifyPrimaryObjects module. To localize iMGs, phase-contrast images were enhanced with the EnhanceOrSuppressFeatures module. Then IdentifyPrimaryObjects segmented individual cells based on size and the thresholding method Robust Background. Measurements were collected using the module MeasureObjectSizeShape and exported to a spreadsheet. (pipeline file: CellProfiler_pipeline_Cells_Area_iMGs.cpproj).

Next we quantified the number and area of synaptosomes present in cells using a similar pipeline applied to the red fluorescence channel images (pipeline file: CellProfiler_pipeline_Red_Area_iMGs.cpproj). First, the GaussianFilter module was used to reduce noise. The images were then thresholded using the RobustBackground method in the Threshold module. Synaptosomes were segmented with the IdentifyPrimaryObjects module, which was set to segment based on both size and the Minimum Cross-Entropy thresholding method. The MeasureObjectSizeShape module recorded size metrics for the synaptosomes, which were then exported to a spreadsheet. Finally, the module OverlayOutlines was used in both pipelines to manually quality check the program’s segmentation of cells and synaptosomes.

